# High-Resolution Additive Manufacturing of a Biodegradable Elastomer with a Low-Cost LCD 3D Printer

**DOI:** 10.1101/2023.06.15.545079

**Authors:** Vahid Karamzadeh, Molly L. Shen, Hossein Ravanbakhsh, Ahmad Sohrabi-Kashani, Houman Savoji, Milica Radisic, David Juncker

**Affiliations:** Biomedical Engineering Department, McGill University, Montreal, QC, Canada; McGill Genome Centre, McGill University, Montreal, QC, Canada; Institute of Biomedical Engineering, Department of Pharmacology and Physiology, Faculty of Medicine, University of Montreal, Montreal, QC, Canada; Research Center, Centre Hospitalier Universitaire Sainte-Justine, Montreal, QC, Canada; Montreal TransMedTech Institute, Montreal, QC, Canada; Institute of Biomaterials and Biomedical Engineering, University of Toronto, Toronto, ON, Canada

**Keywords:** 3D Printing, Elastomers, Tissue engineering, Additive manufacturing, Biomaterials, Photopolymerization

## Abstract

Artificial organs and organs-on-a-chip are of great clinical and scientific interest and have recently been made by additive manufacturing, but depend on, and benefit from, biocompatible, biodegradable, and soft materials. Poly(octamethylene maleate (anhydride) citrate (POMaC) meets these criteria and has gained popularity, and as in principle, it can be photocured and is amenable to vat-photopolymerization (VP) 3D printing, but only low-resolution structures have been produced so fa. Here, we introduce a VP-POMaC ink and demonstrate 3D printing of high resolution (80 µm) and complex 3D structures using low-cost (∼US$300) liquid-crystal display (LCD) printers. The ink includes POMaC, a diluent and porogen additive to reduce viscosity within the range of VP, and a crosslinker to speed up reaction kinetics. The mechanical properties of the cured ink were tuned to match the elastic moduli of different tissues simply by varying the porogen concentration. The biocompatibility was assessed by cell culture which yielded 80% viability and the potential for tissue engineering illustrated with a 3D printed gyroid seeded with cells. VP-POMaC and low-cost LCD printers make the additive manufacturing of high resolution, elastomeric, and biodegradable constructs widely accessible, paving the way for a myriad of applications in tissue engineering, implants, organ-on-a-chip, wearables, and soft robotics.

## 1. Introduction

Organ-on-a-chip platforms integrate microfabrication, tissue engineering, and microfluidics fundamentals to recapitulate relevant aspects of complex living organs *in-vitro*, including microarchitecture, microenvironment, functions of organs, and dynamic cell-to-cell interaction^1, 2^. Organ-on-a-chip engineering could revolutionize new compound screening and biomarker discovery, leading to the development of organ-specific drug screening systems^3^. Fabrication of organs-on-a-chip devices in a biomimetic manner requires great care to achieve tissue-like physiology *in-vitro*. These biofabrication techniques tend to be complex and labor-intensive, and they often require cleanroom facilities which leads to limited growth in the field of organ-on-a-chip. In response to these limitations, 3D printing using bioinks has become an increasingly popular alternative^4^.

3D printing offers many advantages over conventional biofabrication processes for scaffold fabrication, including creating 3D structures with predefined design and fabricating complex 3D tissue constructs similar to biological systems in an automated, rapid, and cost-effective manner^5, 6^. Hydrogels are undoubtedly the most utilized biomaterials for 3D printing and 3D bioprinting due to their properties such as porosity, high water content, and biocompatibility suitable for live cells^7^. However, conventional hydrogels suffer from poor mechanical properties, uncontrollable degradability, and structural stability^8^. Furthermore, to prevent shrinkage and disintegration of the 3D-printed scaffolds, hydrogels must be kept hydrated during and after 3D printing. Additionally, 3D printing must be done in a wet environment to avoid dehydration and shrinkage of the 3D-printed layers.

Driven by the growing need for advanced materials in emerging technologies, research on novel biodegradable elastic polymers has increased, with synthetic, biodegradable elastomeric biopolymers^9, 10^ emerging as promising alternatives to hydrogels due to their tunable and stable mechanical and chemical properties making them suitable for mimicking a wide range of soft biological tissues^11^.

Citrate-based biomaterials have received considerable attention in the last few years due to their biocompatibility, degradability, and flexible designability^12–14^. One of the promising citrate-based biomaterials is poly(octamethylene maleate (anhydride) citrate) (POMaC) elastomer, which is a photo-curable elastomer allowing fast fabrication under mild conditions, degrades through hydrolysis reactions in aqueous solutions and is synthesized from non-toxic monomers^15^. In addition to photocrosslinking of POMaC via the alkene moieties, it can also be thermally crosslinked via easter formation^15^. POMaC has also been successfully used as a key component of implantable pressure/strain sensors for tendons and tissue engineering, selected and proven for its biocompatibility, biodegradability, and mechanical properties. For example, Zhang et al. previously developed an angiogenesis assist device that enabled the merging of two seemingly opposing criteria: permeability and mechanical stability of the vasculature, in a microfabricated polymer-based scaffold for organ-on-a-chip engineering^16^. However, the assembly of the polymer structures involves multiple cumbersome photolithography steps and manual layer-by-layer assembly that hinder the use of POMaC for tissue engineering applications. Photolithography and 3D stamping technology are expensive, lengthy, and involve multistep procedures. 3D printing could provide an alternative solution to such limitations; however, elastomers are particularly challenging to 3D print due to their high viscosity, softness, and slow crosslinking kinetics compared to most common polymers and hydrogels^17^.

To circumvent some of these challenges, 3D printing by extrusion of POMaC followed by photocuring has been explored as an alternative to direct photopatterning; however the high viscosity (>5000 cP) severely limits speed and resolution^18, 19^. Radisic and colleagues developed a coaxial extrusion-based 3D printing of tubular POMaC microstructures with a Pluronic bath^18^. Poly(ethylene glycol) dimethyl ether (PEGDME) was mixed with POMaC to introduce porosity. Thanks to this approach, a variety of perfusable tubes with an inner diameter of 500 µm and a wall thickness of 50 µm could be printed at a much higher speed. However, this method cannot be used for non-tubular structures. In an alternative strategy, Wales et al. used poly(ethylene glycol) diacrylate (PEGDA) with a molecular weight of 700 Da as a copolymer crosslinker with POMaC to improve photoreaction kinetics^19^. POMaC could thus be 3D printed into simple shapes such as rings with a maximal spatial resolution of ∼ 1 mm, which is inadequate for many applications. Additionally, PEGDA had a significant effect on the degradability and mechanical properties of POMaC, which may limit its usefulness in tissue engineering applications. As a result, there is a need for 3D printing of high-resolution and more complex structures made with POMaC.

Vat-photopolymerization (VP) 3D printing is an agglomerative term that encompasses a variety of 3D photopolymerization printing methods, such as stereolithography (SLA) within the vat, which have collectively gained popularity in recent years. These techniques include high-end, expensive 2-photon photopolymerization systems costing > US$500K, widely used digital light processing (DLP, also called stereolithography) systems that typically cost between US$5,000 and US$30,000 with resolutions of up to 2560 × 1600 (∼4M) pixels, and direct laser writers that offer slightly lower resolution and lower throughput but at a somewhat reduced cost of around $2000-$5000. The most common method for VP 3D printing is based on a built platform that progressively moves up as various layers are polymerized by light exposure from the bottom within a vat, forming the 3D print layer-by-layer. However, a back-and-forth movement is normally needed between light exposure and polymerization of the next layer to re-coat the vat with fresh ink prior to its polymerization. Inks with a viscosity lower than 1000 cP are preferred and yield better printability due to more rapid flow and lower suction and layer separation force during the upward movement of the build platform. Most printers are available as light engines with wavelengths of 405 nm, as well as 385 nm and 365 nm, which are often preferred due to the greater availability of inks, higher energy, and increased photo-crosslinking efficiency. More recently, liquid crystal display (LCD) 3D printers have become available at a very low cost of US$300, offering a resolution of 4K (3840 × 2160, ∼ 8M pixels) and, for slightly more expensive models, 8K (7680 × 4320, ∼33M pixels), significantly outperforming more costly DLP printers but only being available with 405 nm illumination. LCD 3D printers have only recently been adopted and explored for research applications^20–22^, but have not been reported for 3D printing of biomaterials.

Herein, we further analyze the challenges that prevented VP 3D printing of POMaC. We then formulate a POMaC ink (VP-POMaC) incorporating cross-linkers, photoabsorbers (PA), diluent, and porogens suitable for printing at a resolution tens-of-micrometers using very low-cost LCD and commercial DLP-based 3D printers. The mechanical properties of prepolymer and polymerized VP-POMaC are characterized, including viscosity, hydrolytic degradation, and compatibility of 3D printed constructs for 3D cell culture. Additionally, we fabricate complex POMaC constructs that were previously unattainable using conventional methods.

## 2. Results

### 2.1. Characterization and Design Criteria for VP 3D Printable POMaC

VP 3D printing of POMaC was challenging because of its low reaction kinetics and its high viscosity (>5000 cP), which make it difficult to produce high-resolution structures, limit printing speed, and the ability to drain uncured ink from the prints, which collectively preclude its use for VP 3D printing. Indeed, a slower reaction rate can lead to decreased resolution because of increased diffusion of the reactants.^23^ It has been reported that a viscosity higher than 3000 cP is not suitable for VP 3D printing, as it would translate to an impractical vat-recoating period between layers of more than 1 min^24^. Additionally, the elasticity and comparatively low mechanical resilience of POMaC can cause damage to the 3D printed structure due to the layer separation force.

In order to 3D print POMaC with an LCD VP 3D printer, we developed the VP-POMaC formulation depicted in **Figure 1** that meets the following criteria. (a) photocrosslinkable with high efficiency at the wavelength of 405 nm available in LCD VP 3D printers, (b) high absorbance at the same wavelength, (c) improved reaction kinetic compared to POMaC, (d) a lower viscosity to facilitate VP 3D printing and draining the uncured ink from microscale features, and (e) suitable biocompatibility for 3D cell culture. These criteria are important to ensure that VP-POMaC could be used effectively with an LCD 3D printer and that the resulting 3D-printed structures would have the desired properties for tissue engineering applications.

**Figure 1.**
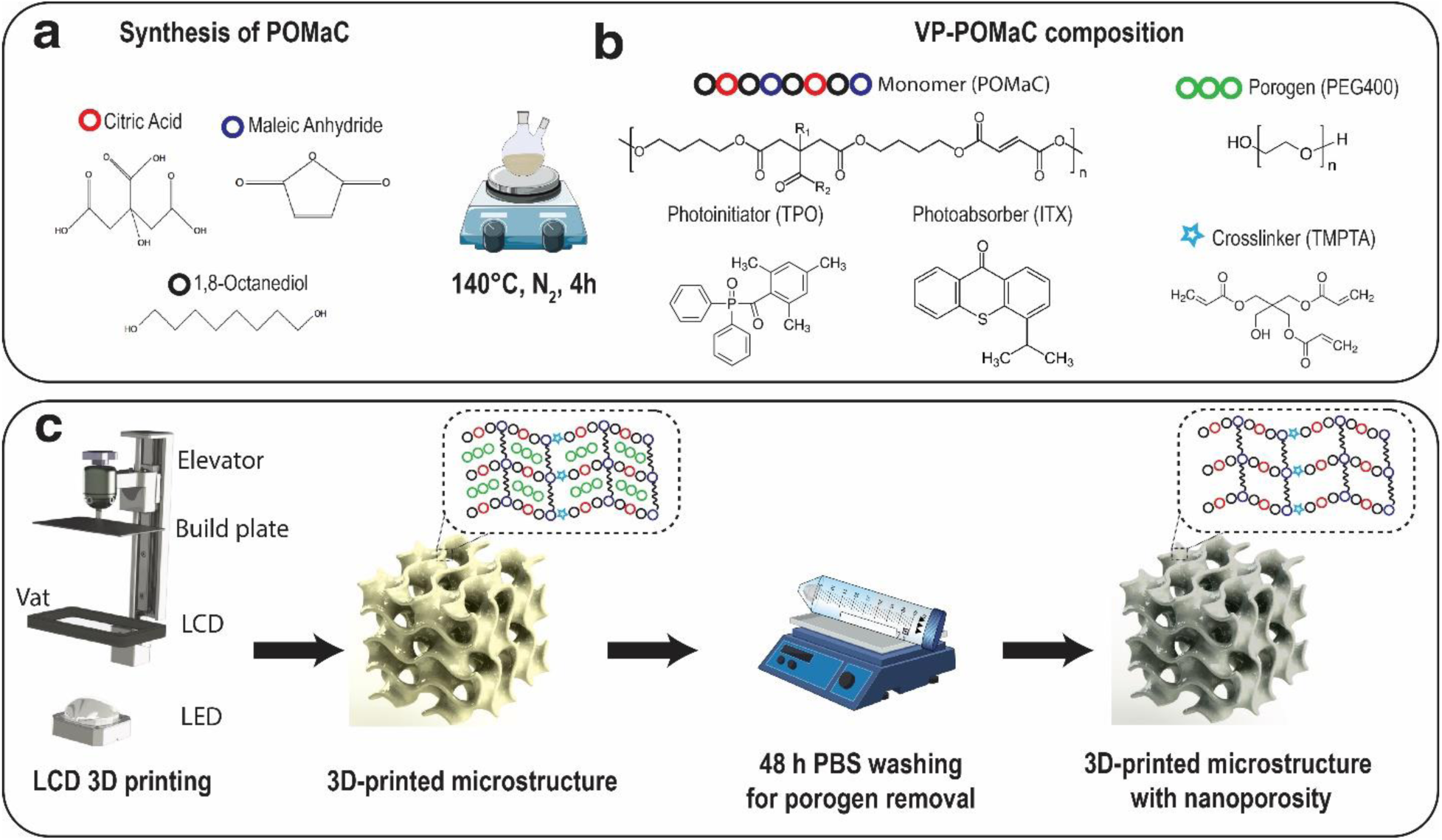
Schematic of the synthesizing and 3D printing process via LCD VP 3D printing. **(a)** POMAC prepolymer was synthesized by mixing three different monomers (citric acid, 1,8-octanediol, and maleic anhydride) at 140◦C for 4 h under N2 purge. **(b)** the VP-POMaC ink composition. **(c)** The fabrication process starts with LCD 3D printing of an object to achieve microporosity, followed by washing steps to remove the porogen to form nanoporosity.

A photoinitiator (PI) is required for VP 3D printing and for our purposes needs to be biocompatible, soluble in POMaC, and has a high absorption at 405 nm. Irgacure 2959 is commonly used as photoinitiator and was used with POMaC, but it has low absorption at wavelengths higher than 375 nm, making it unsuitable for use with 405 nm light. The requirement for sufficient absorbance at 405 nm and low cytotoxicity narrowed down the available photoinitiator options to lithium phenyl-2,4,6-trimethyl benzoyl phosphinate (LAP), 2,4,6- trimethylbenzoyl)-phenylphosphine oxide (BAPO), and diphenyl(2,4,6 trimethylbenzoyl)phosphine oxide (TPO). LAP has the lowest cytotoxicity among these options but is not soluble in POMaC. Compared to BAPO, which is one of the most commonly used photoinitiators in VP 3D printing, TPO is less cytotoxic and has a less yellow color^25, 26^. We thus used TPO, a type 1 photoinitiator that is readily soluble in POMaC at a high concentration (5% w/w). TPO has an absorption wavelength range of 380-425 nm, with a maximum absorption of around 380 nm^27^. While not perfectly matched to the 405 nm light of commercially available 3D LCD printers, it nonetheless remains adequate for efficient polymerization of the POMaC ink. To improve the vertical resolution and prevent clogging of embedded conduits by uncontrolled light penetration through and scattering on cross-linked structures, a photoadsorber is required. We added isopropyl thioxanthone (ITX) as a photoabsorber as it exhibits very low cytotoxicity and is suitable for tissue engineering applications^26, 28^.

Methacrylated/acrylated crosslinkers have been widely employed for free-radical-based polymerization in photosensitive inks. Trimethylolpropane triacrylate (TMPTA) has gained popularity as a crosslinker due to its biocompatibility and the three double-bonds in its chemical structure, which increase the rate of double bonds in the ink and help to improve the reaction kinetics^29^. To facilitate the free-radical-based polymerization of VP POMaC ink, we added 1% (w/w) of TMPTA (912 MW) and found that the gelation time was reduced from 11 s to 7 s. Importantly, TMPTA does not significantly change the mechanical properties of POMaC, so the 3D-printed material retains its desired mechanical behavior even after TMPTA has been added.

The high viscosity of POMaC is an obstacle to 3D printing, and needs to be reduced by either heating in the vat^30, 31^ or by adding a diluent (*i.e.,* a thinner as is used in paint or polish) to the ink^32, 33^. Heating POMaC during the polymerization process could affect its stability and reactivity due to its dual crosslinking characteristic. The diluent is thus the preferred approach but it must be non-reactive and not participate in the crosslinking reaction, while it must leach out after polymerization. For example, reactive diluents such as PEGDA700 help reduce viscosity^19^, but they participate in the crosslinking reaction. Prior works have shown that non-reactive diluents such as PEGDME could also enhance nutrient and oxygen exchange by forming nanopores into POMaC polymer scaffolds^11, 34^. This allows for the production of 3D-printed parts with nanoporosity, which can be beneficial for tissue engineering applications. Hence, we explored the possibility of incorporating polyethylene glycol (PEG) with a molecular weight of 400 a miscible non-reactive material into the ink solution with the expectation that it would reduce viscosity, could be readily leached out after 3D printing thanks to its low molecular weight, and that it would also act as a porogen. PEG is less viscous and expected to be less toxic than PEGDME, which has an ether group. Indeed, PEG is also FDA-approved, water-soluble, and non-toxic and has been used in numerous biomedical and pharmaceutical applications.

By incorporating all the components introduced so far, we developed an optimized ink, VP- POMaC, which was composed of POMaC as the monomer, PEG400 as the porogen and diluent, TMPTA as the crosslinker, TPO as the photoinitiator, and ITX as the PA. We successfully 3D- printed transparent elastomeric microfeatures using VP-POMaC (**Figure 2**). The final construct is obtained after incubating it overnight in 70% ethanol to remove the porogen and unreacted ink components, turning the parts from translucent when freshly printed to their final transparent state. Manufacturing scaffolds with micro-scale features requires high 3D printing resolution. The resolution in VP 3D printing is primarily determined by the projected pixel size and penetration depth of light in XY and Z directions, respectively. However, other factors, such as the ink composition and viscosity, as well as the diffusion of free radicals, can affect the resolution.

**Figure 2.**
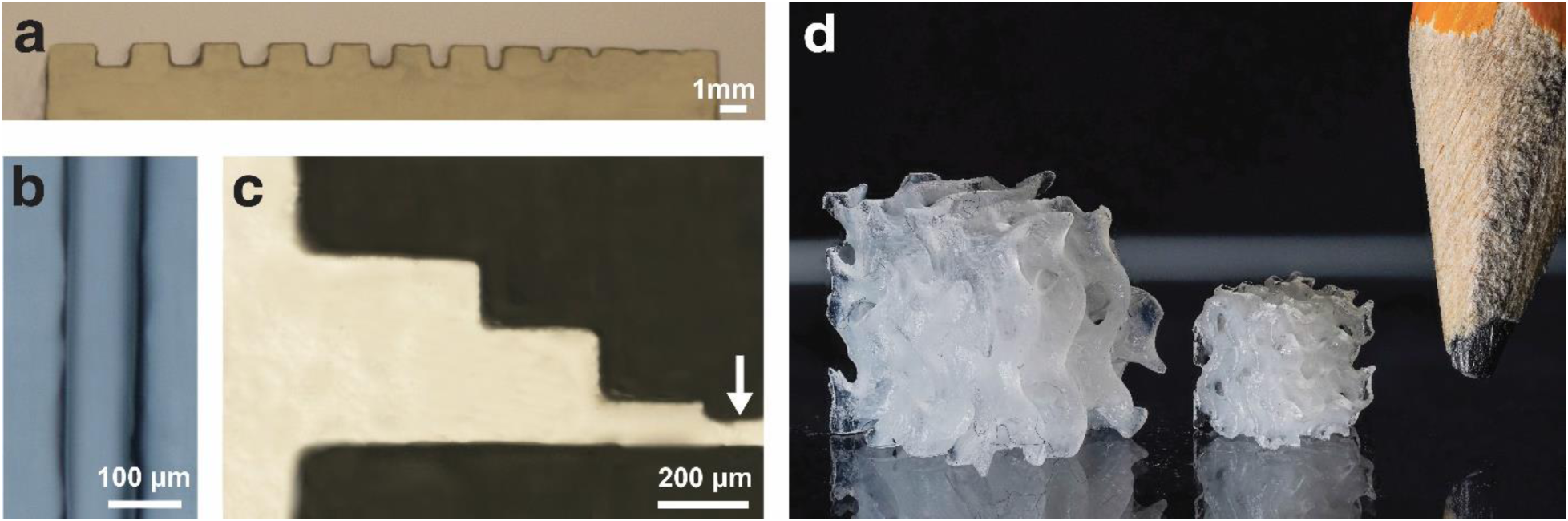
3D constructs made by VP-POMaC ink illustrating its printability. **(a)** Side view of an array of 3D-printed open channels with different dimensions. **(b)** Top-view microscopy image of 3D-printed open channels as small as 100 µm. **(c)** Top-view microscopy image of 3D-printed features as small as 80 µm (white arrow). **(d)** Isometric view of 3D-printed complex gyroid structures with different dimensions. Objects in **(a-c)** were 3D-printed using 40% porogen, while the objects in **(d)** were 3D-printed using 50% porogen.

The LCD 3D printer used in this study has a footprint of 143 × 90 mm^2^ with a projected pixel size of 35 × 35 µm^2^ and 2.5 mw/cm^2^ power intensity. The printing parameters were optimized for producing detailed prints on a footprint approximating the one of a 96-well plate within a few minutes. While DLP-based 3D printers offer sharper pixel edge resolution and higher light intensity, LCD 3D printers are more affordable, have more pixels, and larger printing area. VP- POMaC is also compatible with DLP printers, and 3D structures were printed with shorter exposure times thanks to the higher light intensity of these printers (**Figure S1, Supporting Information**). VP relies on a low adhesion membrane to minimize print failure, and fluorinated ethylene propylene (FEP) membranes are commonly used. Here, to further reduce the separation force, we used a perfluoroalkoxy (PFA) membrane that has a higher tensile strength. This allowed the printed layers to detach more easily and minimized the risk of damaging small features during printing. The layer thickness of 20 µm, in conjunction with a layer exposure time of 50 s, facilitates the production of a smooth surface finish with minimal layer artifacts. After separation in each layer, the resting time of 2 s ensured complete ink recoating during the 3D printing process. Furthermore, the retraction speed was set at a low value of 80 mm/min to reduce the separation force that is proportional to the speed.

In VP 3D printing, the effective Z resolution for overhanging structures is mainly dictated by the penetration depth of light and the reaction kinetics of the ink. The light penetration in the ink can be characterized by measuring the thickness of polymerized ink for different exposure times (**Figure S2, Supporting Information**). Although the exposure time required for curing each layer was reduced from 120 s to 50 s by adding TMPTA, it is still relatively long compared to commonly used non-elastomeric inks. This longer exposure time can lead to the diffusion of free radicals, which can affect the XY resolution of the 3D-printed structures. We evaluated the 3D printing resolution by printing open channels separated by 500 µm gaps with widths ranging from 50 µm to 500 µm in steps of 50 µm. The 50 µm channel was blocked, and the smallest successful 3D- printed channel was measured to be 110 µm, as shown in **Figure 2a-b**. The blockage of the 50 µm channel may have been caused by the scattering of light, the diffusion of free radicals generated during the reaction, or both. Previous research has investigated the potential impact of reactant diffusion and light scattering on the loss of resolution that is commonly observed in VP 3D printing^35, 36^. Additionally, we successfully 3D-printed positive surface features with dimensions as small as 80 µm (**Figure 2c, Figure S3, Supporting Information**). By using the VP-POMaC with 50% porogen and optimized exposure time in our 3D printing process, we successfully fabricated complex structures such as the gyroid with features as small as 100 µm (**Figure 2d**). The gyroid illustrates the capabilities of VP-POMaC and LCD printers for making elastic objects with nanoporosity and intricate microscale geometries that cannot be made by extrusion-based 3D printing or molding of POMaC.

### 2.2. Effect of porogen on printability VP-POMaC and on mechanical properties of 3D- printed parts

We investigated the effect of porogen concentration on the mechanical properties of VP-POMaC using photorheology, compression, and tensile testing. It should be noted that due to the high viscosity of POMaC at concentrations lower than 20% of porogen, preparing samples for tensile testing was challenging, as the elevated viscosity made it difficult to 3D print the material. In contrast, photorheology is compatible with high-viscosity materials, and compression tests necessitate only small disc samples, which can be feasibly produced. As a result, tensile testing was only conducted for porogen concentrations within the 20-50% range, while photorheology and compression testing were conducted for concentrations of 0-50%. Unexpectedly, VP-POMaC with increasing porogen exhibited increasing compression (max. 790 kPa), storage (max. 16.42 kPa), and Young’s modulus (max. 543 kPa) with maximal values found for the maximum porogen concentration of 50% tested in this study. Both compression and photorheology tests confirmed that formulations with a higher concentration of porogen had a higher modulus (**Figure 3a**). Specifically, the compression modulus increased from 80 kPa to 790 kPa as the porogen concentration increased from 0% to 50%. Tensile testing exhibited a similar trend (**Figure 3b**). Additionally, we observed that increasing the porogen concentration resulted in a more brittle material (**Figure 3c and Figure S4, Supporting Information**). Our findings are in agreement with previously reported results on the effect of porogen concentration on the mechanical properties of elastomers^37^.

**Figure 3.**
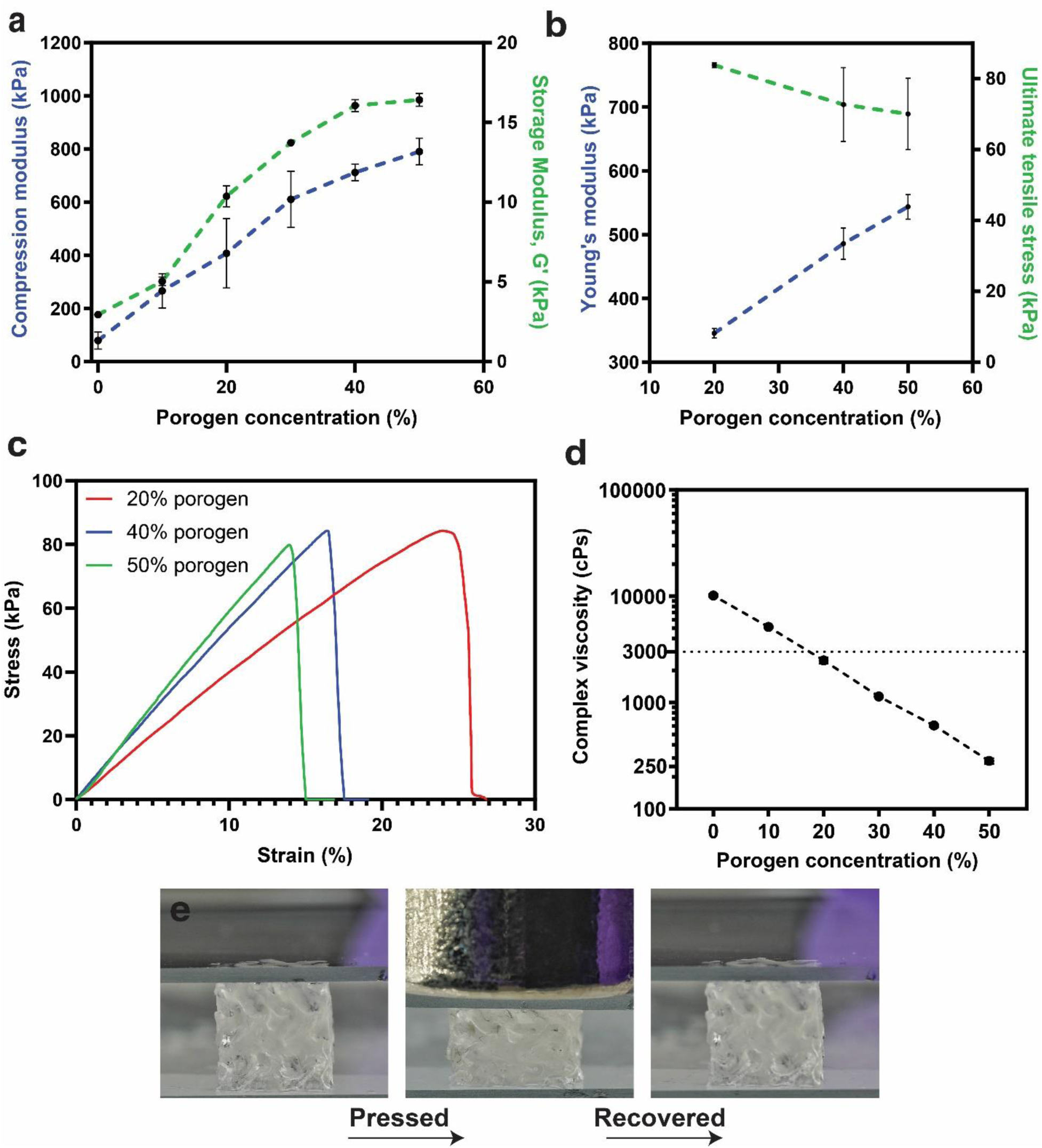
Effect of porogen on the mechanical properties of photocured POMaC. **(a)** Compression and final storage modulus for VP-POMaC with different concentrations of PEG porogen. **(b)** Young’s modulus and ultimate tensile stress for samples 3D-printed with VP-POMaC with different concentrations of porogen. **(c)** Representative stress–strain curves of samples 3D- printed with VP-POMaC prepared with various concentrations of PEG. **(d)** Viscosity for different concentrations of PEG porogen. **(e)**. Images of a 3D-printed gyroid before, while under load, and after recovery **(See Video S1, Supporting Information)**. Error bars represent the standard deviations of three independent experiments (N = 3).

Here, the porogen is primarily utilized as a viscosity-reducing agent to enhance printability. By introducing 50% PEG porogen, the viscosity of POMaC can be reduced from 10136 cP to 282 cP. (**Figure 3d**). We found PEG concentrations of 40% and 50% to be optimal for printing as the reduced viscosity both facilitates that back-and-forth movement of the build plate and the post-printing cleaning, which involves the rinsing and drainage of residual ink, while it also leads to faster photoreaction kinetics.

The increased rigidity within increasing porogen observed in 3D-printed samples may be attributed to several factors. The decreased viscosity of VP-POMaC at higher porogen concentrations may enhance reaction speed due to improved mobility of reactive species. This increased mobility can lead to a higher probability of encounters between reactive species, resulting in a faster curing reaction and ultimately forming a more densely crosslinked network. Moreover, material shrinkage resulting from porogen removal after curing could contribute to an increase in the density of the polymer network. This observation is in agreement with the photorheology and swelling results presented in the subsequent sections. In-depth investigations, beyond the scope of this paper, could offer additional insights into the underlying factors responsible for the observed increase in rigidity. In this study, we focused on the effects of porogen concentrations ranging from 0% to 50% on the mechanical properties of the VP-POMaC. It should be noted that this study did not investigate porogen concentrations higher than 50%; future research could explore the impact of higher porogen concentrations on the material’s properties.

Our findings indicate that the mechanical properties of the VP-POMaC material can be tailored by adjusting the porogen concentration to potentially suit various tissue engineering applications. The storage modulus values, ranging from 2.94 kPa to 16.42 kPa with increasing porogen concentration from 0% to 50%, encompass the desired range for tissues such as skeletal and heart muscle tissues (6-25 kPa)^38–40^. However, it is higher than the range for brain tissue (0.1-1 kPa)^41^ and liver tissue (0.5-3 kPa)^42^.

### 2.3. Effect of porogen on the crosslinking rate

To investigate the impact of PEG porogen and TMPTA on POMaC’s gelation time, we conducted photorheology tests to measure the storage modulus (G’) and loss modulus (G’’) during UV light exposure. The gelation time was determined by the point at which G’ crossed over G’’, indicating the material’s transition from a gel to a solid state. Before crosslinking induced by UV light (<15 s), the G’ and G’’ values were stable (**Figure 4a**). The addition of 1% TMPTA significantly reduced the gelation time from 11 s to 7 s in the photorheometer, illustrating the influence of low concentrations of the TMPTA crosslinker with three double bonds on reaction kinetics (**Figure 4a and Figure S5, Supporting Information)**. This has notable benefits for 3D printing, where rapid polymer crosslinking during printing is desirable. The effect of porogen concentration on the crosslinking rate of the polymer is also considerable, as illustrated in **Figure 4b**. Increasing the porogen content exhibited a clear acceleration in crosslinking rate. This may be attributed to the significantly lower viscosity of the VP-POMaC ink at higher porogen concentrations (**Figure 3d**), which enables the greater potential for the increased free radical diffusion coefficient, resulting in a more pronounced impact on the crosslinking rate. The same underlying mechanism could explain the correlation observed between mechanical properties and porogen concentration.

**Figure 4.**
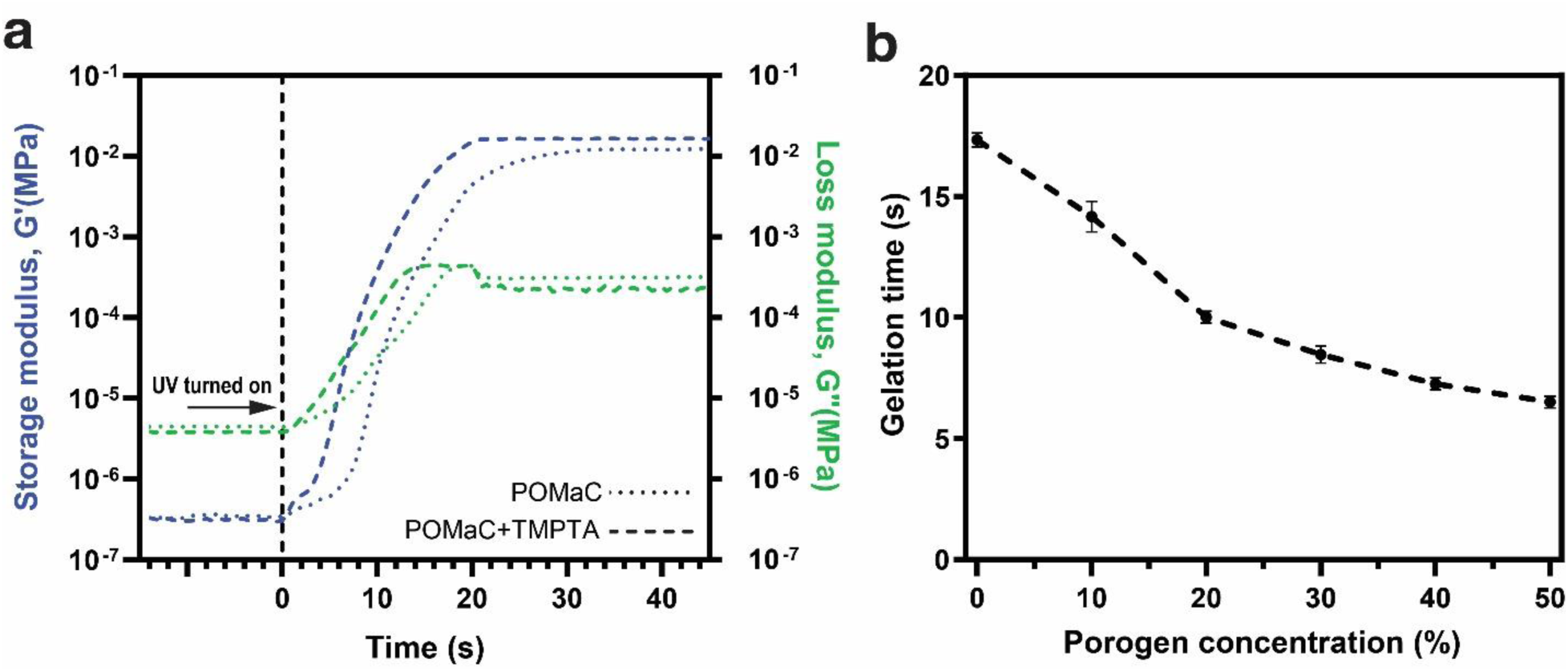
Crosslinking rate, gelation time of POMaC and VP-POMaC. **(a)** time sweep photorheology measurements of POMaC and VP-POMaC ink both with 40% porogen. The addition of TMPTA in the VP-POMaC ink formulation resulted in a decrease in gelation time from 11 s to 7 s as compared to the ink formulation without TMPTA. **(b)** Gelation time under UV illumination for different concentrations of porogen. The gelation time is reduced when the concentration of porogen is increased.

### 2.4. *In vitro* swelling and degradation

Bioinks with low swelling/ shrinkage ratios (<50%) are favorable for tissue engineering and wound healing applications^43^ as a low swelling ratio is essential for maintaining the architecture and fidelity of 3D-printed microstructures under physiological conditions. The swelling behavior of VP-POMaC inks was characterized by incubating them in PBS for up to 21 d. The shrinkage increased for samples with higher concentrations of porogen (**Figure 5a and Figure S6, Supporting Information)**. This could be ascribed to the higher porosity of samples with porogen, which leads to higher water uptake. Notably, the shrinkage for all concentrations remained below 25%, indicating a low level that is suitable for tissue engineering applications.

**Figure 5.**
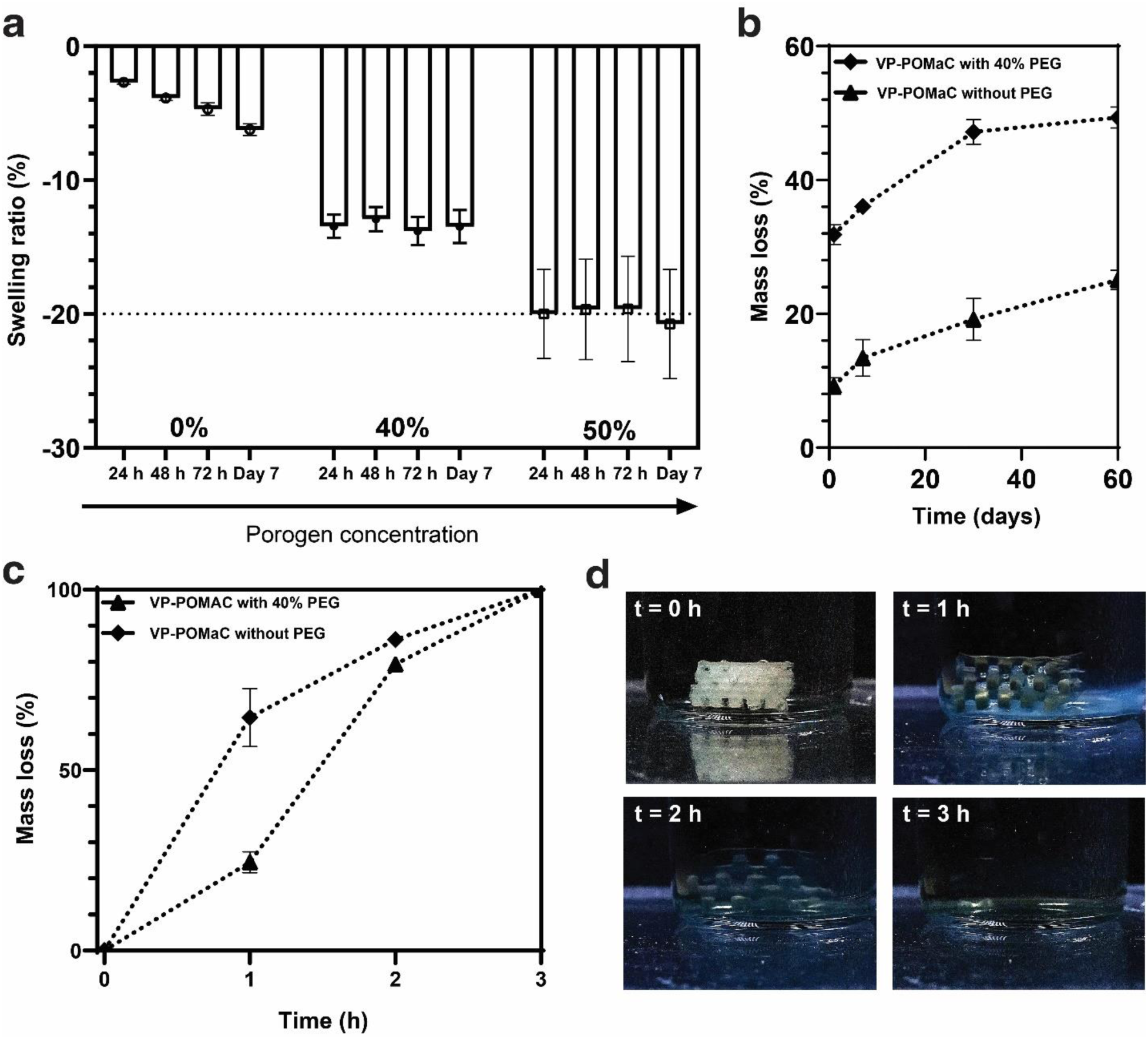
*In-vitro* swelling and degradation of VP-POMaC. **(a)** Swelling ratio over a week in PBS solution. **(b)** mass loss of VP-POMaC disks without porogen and with 40% porogen in PBS solution at 37°C over 60 days and **(c)** accelerated mass loss in 0.25M NaOH solution Representative of three independent experiments (N = 3). **(d)** hydrolytic degradation via surface erosion of 3D printed woodpile structure with 40% porogen over time in 0.25 M NaOH solution **(See Video S2, Supporting Information)**.

Biodegradability is essential for tissue-engineered scaffolds, and the rate of degradation requires to match the tissue formation rate, to finally replace the regenerated tissue. Hence, the long-term performance of scaffolds *in-vivo* is strongly dependent on the degradation rate. The polymer molecular structure and composition have a critical impact on the degradation rate. The ester bonds in the POMaC backbone are hydrolytically degradable, and depending on their relative content, they mediate controlled degradation rates^37, 44^. When exposed to physiological conditions, POMaC degrades via surface erosion, a process that sequentially breaks the ester bonds between monomers.

To investigate the *in-vitro* degradation of VP-POMaC, we fabricated disks via 3D printing and incubated them in phosphate-buffered saline (PBS) buffer (pH 7.4) and sodium hydroxide (NaOH) at 37° C. As shown in **Figure 5b**, the samples containing 40% porogen experienced significant weight loss in PBS compared to pure VP-POMaC. The rate of degradation significantly slowed down after day 1, and samples with PEG porogen exhibited 50% mass loss after 60 days. The initial mass loss (∼10%) is associated with soluble low molecular weight chains in the polymer, while the additional mass loss in the porous VP-POMaC samples is due to the leaching of water-soluble PEG porogen.

Polymers that contain ester linkages, such as POMaC, are more susceptible to hydrolytic degradation via surface erosion in the presence of NaOH, which hydrolyses the ester groups and can speed up degradation. We investigated the accelerated degradation of POMaC at a concentration of 0.25 M sodium NaOH solution. 3D-printed structures were immersed in 0.25 M NaOH solution at room temperature with no agitation. The VP-POMaC samples were completely degraded and dissolved after 3 h in an aqueous base solution (**Figure 5c, Supplementary Video S2**). **Figure 5d** shows the dissolution as time progresses. Most of the photocurable inks are derived from acrylates and epoxides that are difficult to degrade once crosslinked. 3D printable materials such as POMaC can address the concerns about pollution issues for VP inks that are of increasing concern as 3D printing becomes more widely adopted.

### 2.5. Cytocompatibility and 3D cell culture

POMaC has been widely used for cell-based applications which require a high level of biocompatibility. As unreacted POMaC ink components remain in the 3D-printed part and are cytotoxic, extensive post-printing washing is needed to eliminate all unreacted residues prior to cell culture. The washing step is also necessary for the pore formation via extraction of the porogen component. High porosity could be favorable for applications with cells, as it may facilitate cell adhesion, sprouting, and exchange of nutrients^16, 45^. To achieve optimal biocompatibility and porosity, 3D-printed POMaC samples were washed with PBS and 70% EtOH for at least 48 h to remove any residual porogen and unreacted ink components. Throughout the washing process, we alternated the wash buffer between PBS and 70% EtOH every 12 h to best utilize POMaC’s distinct swelling behavior in each buffer.

VP-POMaC was tested for cytocompatibility using the human lung fibroblast cell line (IMR-90) and human umbilical vein endothelial cells (HUVEC). The cell lines were chosen for their wide usage in research, as well as for their intolerance of sub-optimal culturing conditions. The cytocompatibility assays were performed in compliance with International Organization for Standardization (ISO) standards (10993-5:2009) for the development of medical devices. According to ISO 10993-5:2009 standard, the 3D-printed POMaC samples were post-treated with the above-mentioned washing process and directly co-cultured with our model cell lines in a 96- well plate. In addition to a cell-only control condition, a previously developed POMaC formulation^34^ was used as a cytocompatibility benchmark. Microscopy images and quantitative cell viability measurements via PrestoBlue™ were taken every 24 h for a time course of 3 days. We observed that HUVECs and IMR-90 co-cultured with VP-POMaC or the benchmark POMaC formulation were morphologically indistinguishable compared to the cell-only control condition **(Figure 6 and Figure S7, Supporting Information)**. Furthermore, the quantified cell viability (normalized to cell-only control) showed excellent biocompatibility (>80% for all time points) of both VP-POMaC and the benchmark POMaC formulation.

**Figure 6.**
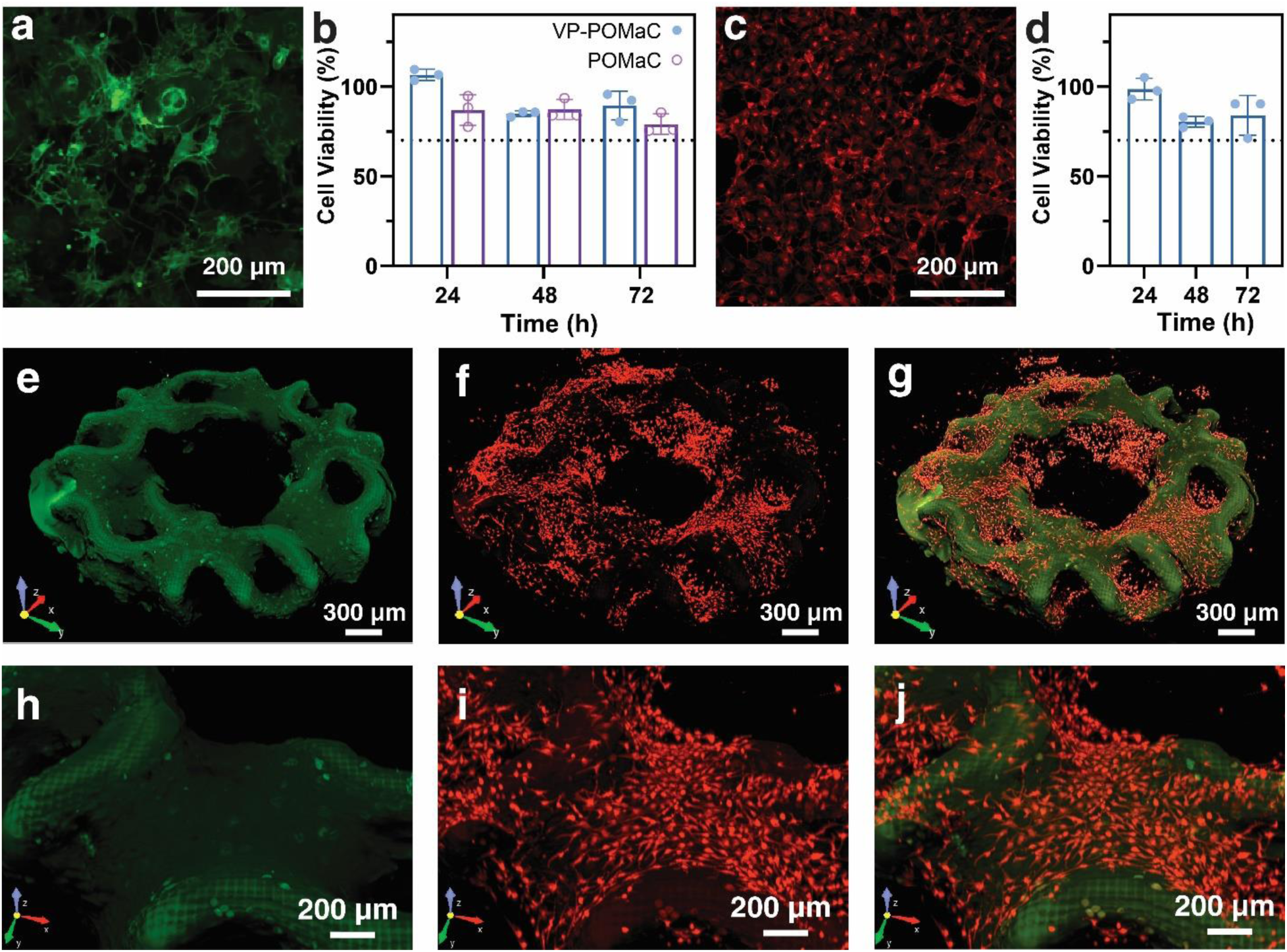
3D cell culture on VP-POMaC scaffold. **(a)** Fluorescence images of GFP labeled IMR- 90 human lung fibroblast cells following 48 h culture in a well-plate in presence of VP-POMaC. **(b)** IMR-90 cell viability when cultured with VP-POMaC or POMaC for up to 3 d. **(c)** Fluorescence image of mCherry-labelled HUVEC cells following 48 h culture in a well-plate in the presence of VP-POMaC. **(d)** HUVEC Cell viability for 3 d. **(e-g)** 3D confocal microscopy images of HUVEC cells (red) cultured and adhered on an autofluorescent cylindrical gyroid scaffold (green) along with close-up images **(h-i)** A close-view of cultured HUVEC cells (red) reveals cellular proliferation on the scaffold (green).

In addition, we cultured HUVEC cells on a 3D-printed cylindrical lattice gyroid structure as a demonstration of cell-populated 3D-printed POMaC structures for prospective tissue engineering applications. The scaffold was designed with voids with a minimum diameter of 200 µm. To enable cell attachment, the 3D-printed VP-POMaC structure was incubated in cell culture media supplemented with 40 µg/mL collagen I overnight at 4℃. The morphological characterization shows a satisfactory cellular adhesion to the scaffold, as illustrated in **Figure 6d-f**. In 3D culture, cells were found to proliferate throughout the entire scaffold for pore sizes between 200-400 µm. The scaffold exhibits autofluorescence in the green channel, which aids in the visualization of its topography and cell attachment. In summary, VP-POMaC ink shows good biocompatibility, cell adhesion, and proliferation, making it a promising candidate for various tissue engineering applications.

## 3. Discussion and Conclusion

3D printing has transformed tissue engineering, offering unprecedented freedom in designing and fabricating intricate microstructures that are challenging to create using traditional fabrication methods. The use of elastic, biodegradable materials is desirable for many tissue engineering applications, and POMaC has emerged as a candidate material but has been difficult to shape by 3D printing. Here, we introduced VP-POMaC and the use of DLP and low-cost LCD printers for the fabrication of complex scaffolds that were unattainable with previous fabrication methods for POMaC.

This presents a notable breakthrough in the 3D printing of POMaC, achieving a resolution down to tens of microns with the smallest feature printed at 80 µm. The optimized exposure time of 30 s to 50 s per 20 μm layer constitutes a major improvement over the 4 min crosslinking time (per layer) required in previously reported manual stamping processes, which also entailed labor-intensive manual alignment and multiple masks for each scaffold, ultimately extending the total scaffold fabrication time to a day. The DM process for POMaC 3D printing is largely automated, eliminating the need for expensive equipment and manual labor while also enabling faster design iteration. In addition to the faster fabrication time, using this approach allows for the fabrication of more complex scaffolds, such as gyroid structures that were previously impossible to create. These complex scaffolds can be utilized in various tissue engineering and organ-on-a-chip (OoC) applications that demand precise designs.

By reducing the gelation time from 11 s to 7 s as measured by photorheology, adding a crosslinker with three double bonds, and decreasing the viscosity using PEG porogen, we successfully adapted POMaC for VP printing on the low-cost LCD 3D printer. The resulting POMaC constructs exhibit high biocompatibility (>80% cell viability) with fibroblast and endothelial cells. Their mechanical properties can be tailored to suit specific tissue engineering needs with a range of storage moduli (2.94-16.4 kPa) and compression moduli (79.3 - 790 kPa) suitable for various applications, such as skeletal and heart muscle tissues. This controllability of mechanical properties offers the potential to create biomaterials that more accurately recapitulate the tissue-specific biomechanical microenvironments, thereby enhancing the prospective efficacy of tissue engineering approaches. Additionally, this flexibility in the mechanical profile of the material broadens its application spectrum in non-biomedical fields. Moreover, compatibility with both DLP and low-cost 3D LCD printers (available for under $300) will open up new opportunities for VP-POMaC, tissue engineering, and VP printing,

Future work will be needed to investigate the effect of PEG porogen on the pore size, resolve the low printability at lower porogen concentrations, and validate or improve the bioink for high-resolution 3D printing down to the sub-ten-micrometer scale necessary for reproducing small embedded vasculature networks that correspond to the size range of capillaries. Beyond tissue engineering, the 3D-printed POMaC constructs also hold promise for use in wearables, soft robotics and possibly flexible electronics thanks to its biocompatibility and tunable elastomeric properties. Hence, the convenient and rapid fabrication of biodegradable, biocompatible, and elastic microstructured POMaC constructs using an affordable LCD 3D printer creates a wealth of opportunities not just for tissue engineering, implants, and organ-on-a-chip, but also for wearables and soft robotics.

## 4. Materials and Methods

### 4.1. 3D printing

All objects were designed in SolidWorks, exported as “STL” files, and 3D-printed with a ELEGOO Mars 3 LCD 3D printer (ELEGOO, China) with a 35 µm pixel size and 4K monochrome LCD. All objects reported in this work are printed with a layer thickness of 20 µm and an exposure time of 50 s at a light intensity of 2.2 mW/cm^2^. Immediately after printing, to remove unpolymerized ink, objects were washed with 70% EtOH (Fisher Scientific, Saint-Laurent, Quebec, Canada) several times and dried under a stream of pressurized nitrogen gas. Then, the objects were immersed in 70% PBS for 48 h to remove the residual ink and porogen.

### 4.2. Synthesis of POMaC

Poly(octamethylene maleate (anhydride) citrate) (POMaC) was synthesized as described previously^46^. Briefly, maleic anhydride, citric acid, and 1,8-octanediol were mixed in a two-necked round bottom flask at a 2:3:5 molar ratio and melted at 160 ℃ under nitrogen purge, and the mixture was stirred for 2 h. The resultant prepolymer was then dissolved in 10 ml 1,6-dioxane and purified through drop-wise precipitation in deionized distilled water. Then the collected polymer was concentrated and dried under airflow for two days and stored at 4℃. Following drying, the prepolymer was analyzed using FT-IR spectroscopy (**Figure S8, Supporting Information**), and the obtained data was consistent with the spectra reported in prior studies^15, 19^.

### 4.3. Formulation of VP-POMaC ink

Purified POMaC was mixed with a PEG (no. 202398; Sigma-Aldrich), which is miscible with both POMaC and water and has a low molecular weight, allowing it to be leached out after crosslinking of the polymer. Different concentrations of PEG, spanning from 10% (w/w) to 50% (w/w), were used to quantify the effect of porogen on the mechanical and reaction kinetic characteristics of the ink. TMPTA (no. 412198; Sigma-Aldrich) at a concentration of 1% (w/w) was added to the ink to decrease the onset time by providing more acrylate groups.

The PI, TPO (no. 415952; Sigma Aldrich), was mixed with the POMaC/PEG solution at a 2 % (w/w) concentration after preliminary experiments were conducted to identify a concentration that produced prints of adequate resolution (i.e., lower concentrations did not consistently produce prints with a satisfactory resolution, while higher concentrations did not necessarily cause any improvement). To assist in mixing the PI into the solution, the solution was heated on a hot plate at 60-90 °C for 2-5 min and then mixed for 30-120 min, depending on the mixing progress. The other component of the ink was a PA, ITX (no. TCI0678; VWR International) was mixed with the ink at a concentration of 0.8% (w/w).

### 4.4. Videos and image stacking

Videos and images were captured using a Panasonic Lumix DMC-GH3K and Sony α7R III. Focus stacking was performed using Imaging Edge Desktop (Sony Imaging Products & Solutions Inc., Japan) to obtain a sequence of images at various focal planes. The images were then processed using CombineZP (available at https://combinezp.software.informer.com/). Microscale imaging was conducted on a Nikon Eclipse LV100ND inspection microscope at 5x magnification. Some images were captured using a Leica SMZ-8 stereo microscope equipped with a Lumix GH3 DSLR digital camera from Panasonic.

### 4.5. Photorheology

A Discovery HR-2 rheometer (TA-Instruments, DE, USA) was used in conjunction with the ultra- violet (UV) accessories to investigate the effect of porogen and TMPTA on the storage and loss moduli and the gelation time of the ink. A volume of 200 μL was added to the bottom plate of the rheometer, and a time sweep test was performed while the sample was exposed to UV radiation for 45 s. The UV light intensity was set at 20 mW.cm^-2^. The time sweep test was conducted at room temperature using a torsional frequency of 1 Hz and a torsional strain of 1%. The experiment was performed for different study groups, and the storage and loss moduli were recorded to find the onset time as well as the gelation time, i.e., the intersection of the loss and storage moduli curves.

### 4.6. Compression and tensile test

Disk-shaped samples were 3D-printed with a diameter of 10mm. The samples were then washed in 70% EtOH (Fisher Scientific, Saint-Laurent, Quebec, Canada) for 48 h before the compression test. A dynamic mechanical analyzer (DMA, Q800 TA-Instruments, DE, USA) was employed to quantify the compression modulus of the crosslinked samples. Tensile testing was carried out using an Instron 3360 electronic universal testing machine (Instron corporation, MA, USA).

### 4.7. Swelling/shrinkage

Disk-shaped samples were prepared similarly to samples for compression tests. All the samples were weighted after crosslinking. The samples were then soaked in PBS 1x and shaken over an orbital shaker for certain time periods of up to 3 weeks. At each time point, the samples were removed from PBS, weighed, and resoaked in PBS for the next time point. The solution was changed at every time point to avoid saturation.

### 4.8. *In-vitro* degradation

Three disk-shaped samples, each with a diameter of 8 mm and a thickness of 3 mm, were fabricated for performing degradation in PBS 1x and 1M NaOH at 37°C. After crosslinking, the samples were frozen at -80°C for 24 h and then lyophilized for 72 h using a ModulyoD 5L freeze dryer (Thermo Fisher Scientific) at 290 ± 10 μbar and room temperature. The samples were then weighed and soaked in PBS 1x or 1M NaOH for different periods up to 8 weeks and 3 h, respectively. At each time point, a set of samples was collected, and the media was changed for other samples. At the final point, all samples were frozen, lyophilized, and weighed to find the remaining weight fraction.

### 4.9. Penetration depth measurements

In order to measure the penetration depth of light, a drop of the formulation was placed on a glass slide and exposed to different exposure times at a light intensity of 2.2 mW/cm^2^. After rinsing uncured ink with 70% EtOH, we measured the thickness of patterned regions using a stylus profilometer (DektakXT, Bruker Co.).

### 4.10. Cell culture

The normal human fibroblast cell line IMR-90 (ATCC CCL-186) expressing the fluorescent protein GFP was cultured in Dulbecco’s Modified Eagle Medium (Gibco, USA) containing 4.5 g/L D-glucose, 4.5 g/L L-glutamine, and 110 mg/mL sodium pyruvate. The media was further supplemented with 10% FBS and 1% penicillin-streptomycin (Gibco, USA). All cell lines were incubated at 37°C with 5% CO2 supplementation. Human umbilical vein endothelial cells (HUVEC) expressing the fluorescent protein mCherry were kindly provided by Dr. Arnold Hayer of McGill University^47^. HUVEC cells were cultured in EGM-2 media (Lonza, USA). The cells were grown and passaged according to ATCC’s recommendations.

### 4.11. Cytotoxicity Testing

The cytocompatibility assays were performed in compliance with International Organization for Standardization (ISO) standards (10993-5:2009) for the development of medical devices. POMaC disks with a thickness of 800 µm were made by UV curing 150 μL of VP-POMaC ink in a PDMS-coated 24-well plate. The disks were washed for at least 72 h in PBS and EtOH to leach out any unreacted ink, as well as the porogen. IMR-90 cells were seeded in a 24-well plate at a seeding density of 10k per well. Immediately after cell seeding, post-treated POMaC disks were placed directly on top of the cell layer. As controls, cell-only wells, as well as cells co-cultured the literature POMaC formulation, were seeded with the same cell density. Quantitative cell viability measurements via PrestoBlue™ (Thermofisher, USA) were taken every 24 h for a time course of three days. Microscopy images were taken daily using a Ti2 inverted microscope and analyzed using NIS-Element (Nikon, Japan). Three biological replicates were done for each of the three above-mentioned conditions. For the PrestoBlue™ cell viability assay measurement, two technical replicates were performed for each biological repeat.

### 4.12. 3D co-culture of POMaC and HUVEC

3D-printed POMaC structures were post-treated as described above. Additionally, they were incubated in cell culture media supplemented with 40 µg/mL collagen I overnight at 4 degrees Celsius to enable cell attachment to the surface of the structure. Trypsinized HUVEC cells were washed once in PBS, resuspended in EGM-2 media at a concentration of 1x10^6^ cells/mL, and supplemented with 10% growth factor reduced Matrigel^®^ (Corning, USA). HUVECs were seeded in droplets onto the air-dried POMaC structure and incubated in the cell culture incubator for 1 h to allow cell attachment. Afterward, the POMaC structure was flipped upside-down, and the seeding was repeated for the opposite side. Finally, the co-culture was carefully transferred into an ultra-low attachment 96-well microplate (Corning, USA) and cultured in EGM-2 media for at least 24 h prior to confocal imaging.

### 4.13. Fourier Transform Infrared (FT-IR)

To evaluate the synthesizing process, FTIR (Thermo-Scientific Nicolet 6700, Waltham, MA, USA) was conducted on the prepolymer and VP-POMaC (**Figure S8, Supporting Information**).

### 4.14. Data analysis

The error bars displayed in the figures represent the standard deviation. GraphPad Prism 9 was used for data analysis.

## 5. Contributions

V.K. developed the VP-POMaC formulation, designed and performed experiments, analysed data and prepared the manuscript. M.S. carried out the cytotoxicity assay, and 3D cell culture. H.R. contributed to the mechanical testing and polymer synthesis. A.S. contrubioted to the swelling assesment. H.S. provided advice and trained V.K. on the POMaC synthesis process. V.K., M.R., and D.J. conceived the idea. D.J. provided funding, supervised the project, and contributed to writing the original draft, reviewing, and editing the manuscript.

## 6. Acknowledgments

Authors acknowledge Y. Paschalidis, V. Han, S. Okhovatian, S. Campbell and P. Chimienti for their assistance and Y. Morocz for photography. V.K. acknowledges FRQNT for the doctoral scholarship (no. 268838). D.J. and M.R acknowledge support from NSERC strategic grant (no. 506689-17). M.R. acknowledges support from a Canada Research Chair in Organ-on-a-Chip Engineering. D.J. acknowledges support from a Canada Research Chair in Bioengineering.

## Supplementary Information

1. 3D-printed parts with DLP

2. Penetration depth

3. 3D-printed Angiochip scaffold

4. Mechanical testing

5. Photorheology

6. Swelling

7. Cytotoxicity

8. FTIR

## Supplementary Videos

**Video S1.** Flexible gyroid structure

A flexible gyroid structure 3D-printed with VP-POMaC (40%) under load without a change in the shape.

**Video S2.** VP-POMaC degradation in NaOH

Rapid degradation of VP-POMaC at different NaOH concentrations

**Figure S1.**
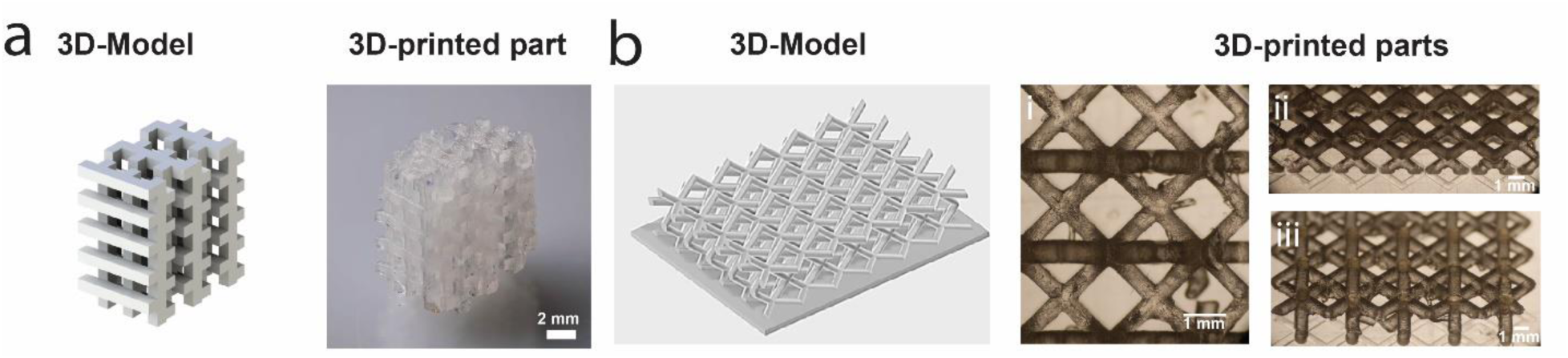
3D VP-POMAC constructs made with a with DLP 3D printer. **(a)** woodpile model and 3D print**. (b)** Scaffold with freestanding rods including the model and the prints. Microscopy images of **(i)** top-view, **(ii)** and **(iii)** side-views of 3D-printed part.

**Figure S2.**
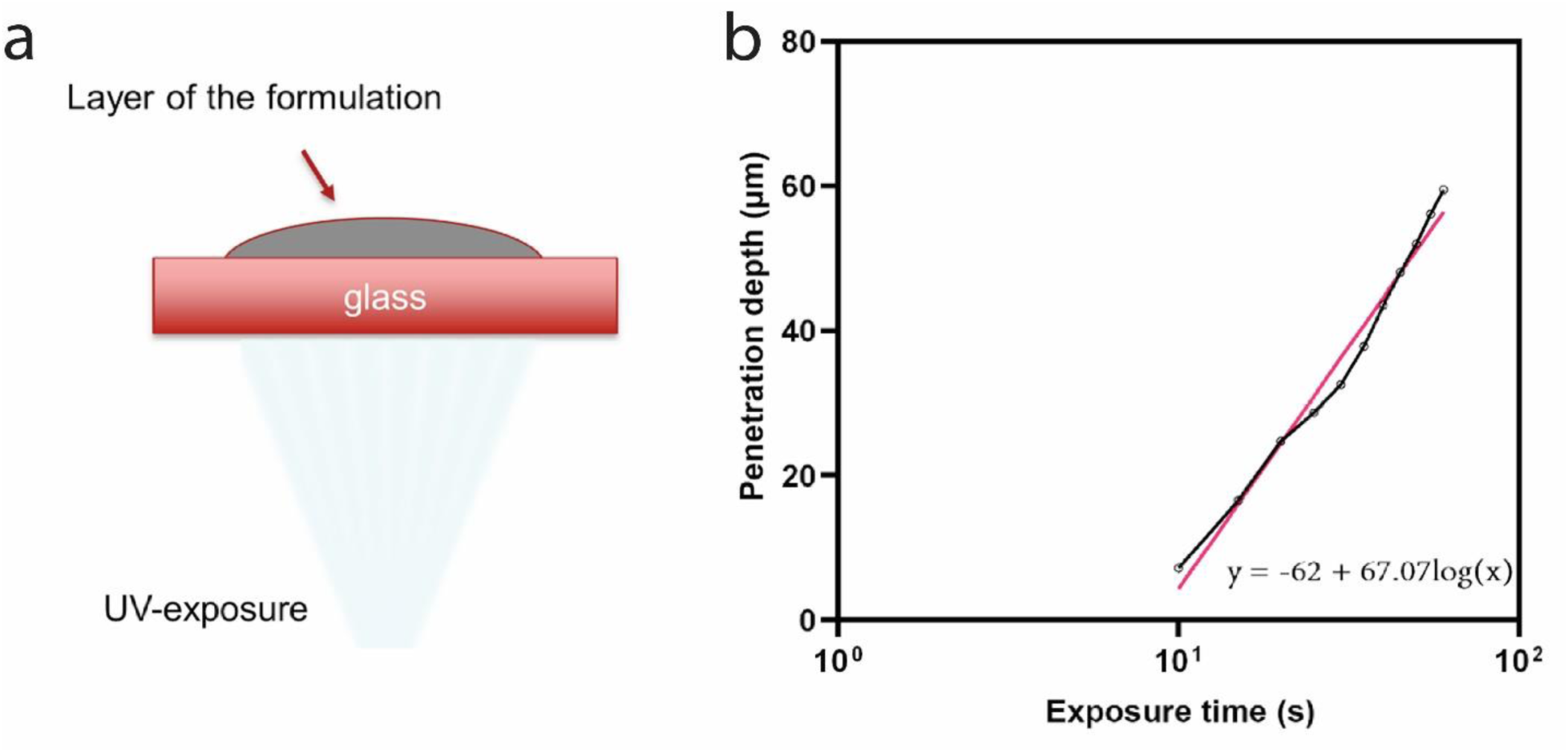
No text of specified style in document.. Penetration depth characterization of the VP-POMaC with 40% PEG porogen. **(a)** The image shows the setup with UV light at the bottom, glass, and VP-POMaC ink on top. Following UV exposure, uncured VP-POMaC was removed by rinsing. **(b)** The graph shows the thickness of polymerized PoMAC measured by a profilometer as function of the exposure time thus reflecting the penetration depth of the UV light.

**Figure S3.**
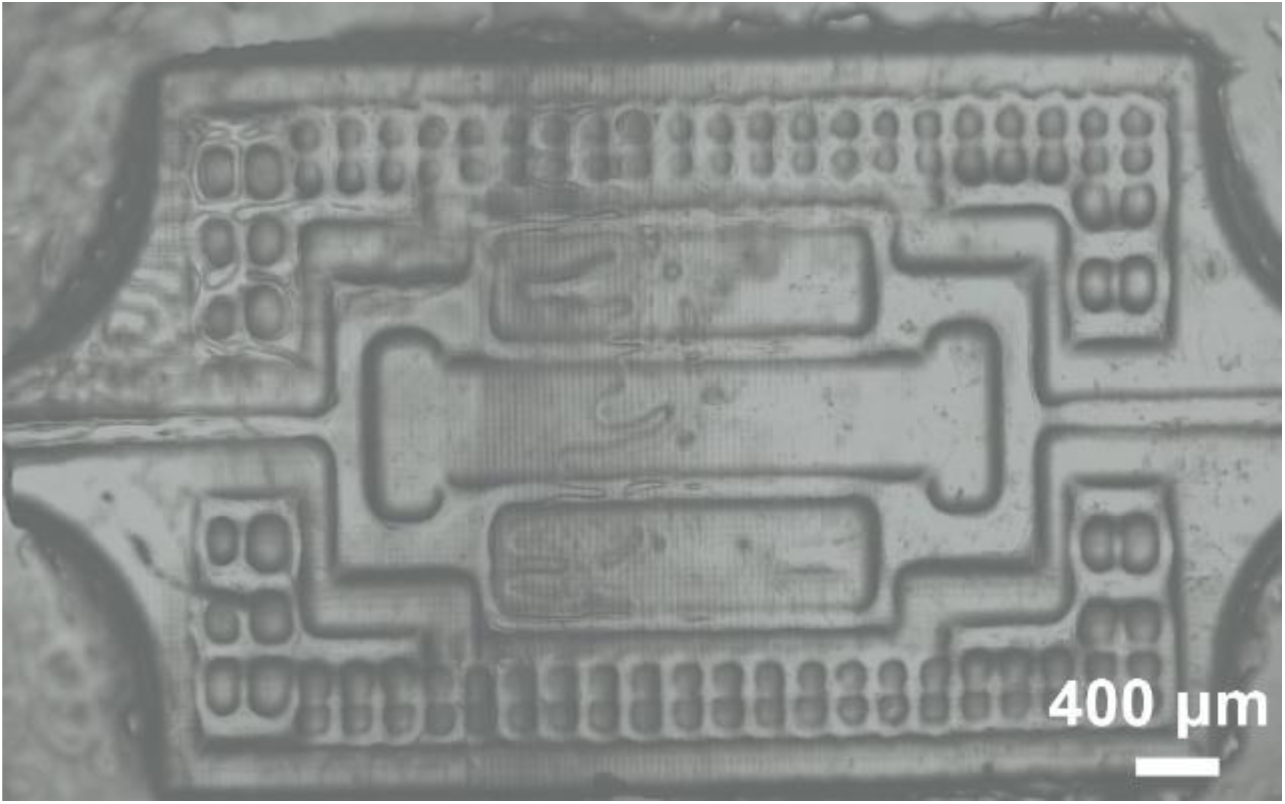
No text of specified style in document.**. 3D-printed Angiochip scaffold with open channels.** The smallest features in this scaffold are 150 µm in size.

**Figure S4.**
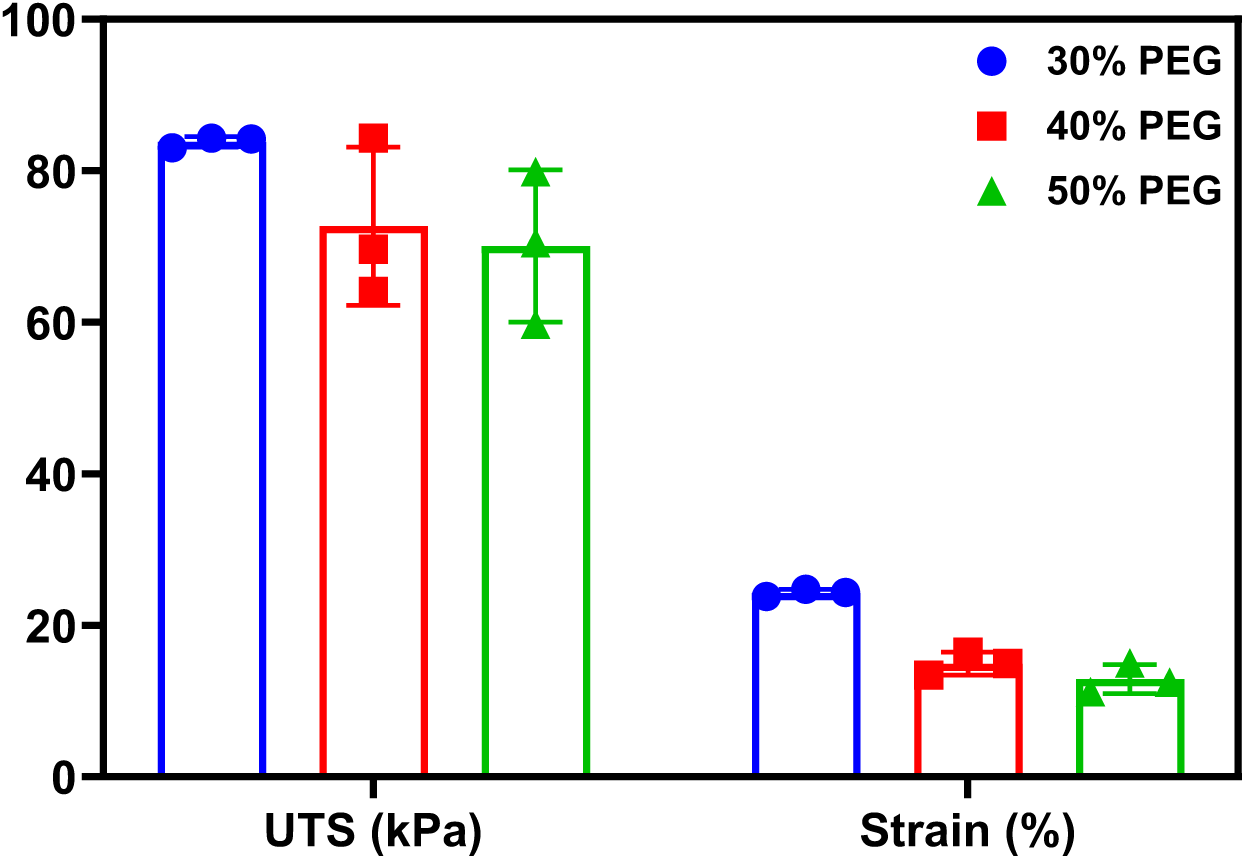
No text of specified style in document.. Ultimate tensile stress (UTS) and maximum strain of VP-POMaC with different concentrations of porogen. Both UTS and maximum strain exhibited a decrease with higher porogen concentrations.

**Figure S5.**
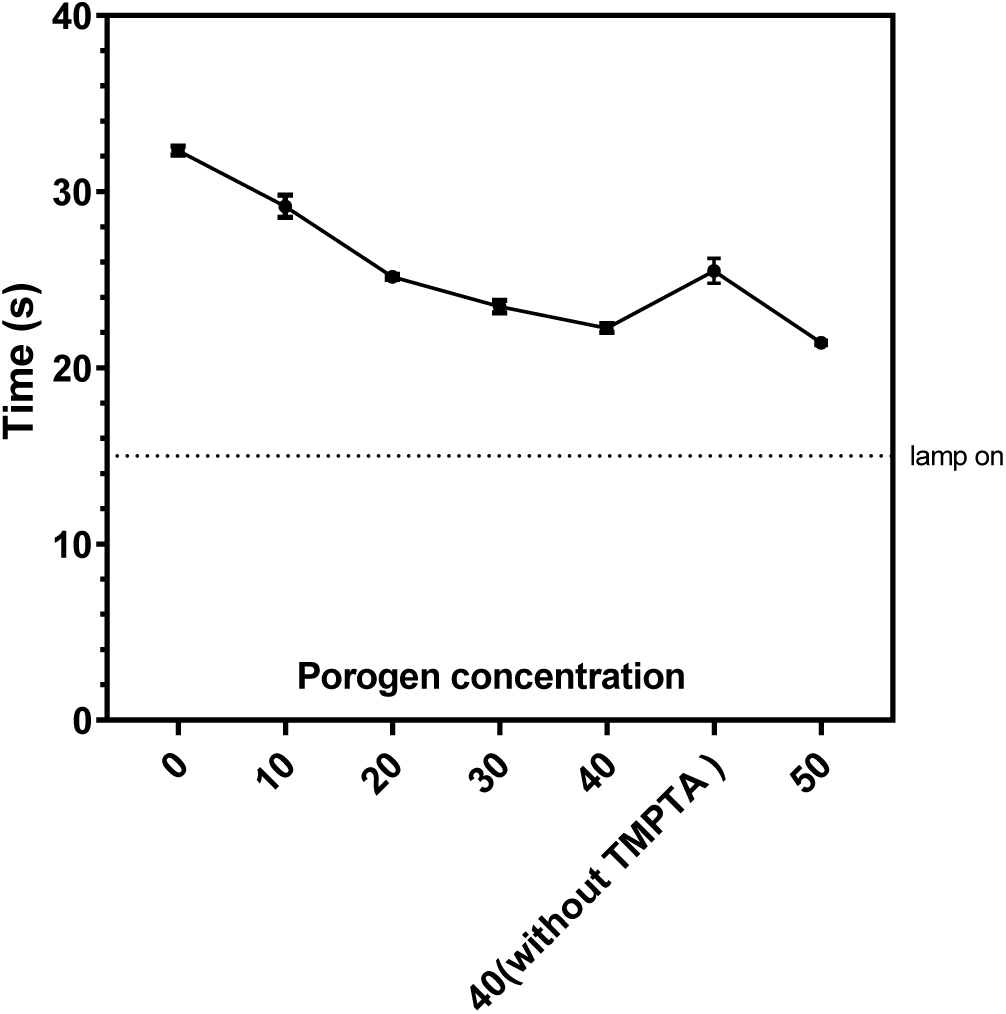
Effect of porogen and TMPTA on the gelation time. In common VP-POMaC formulations with TMPTA, increasing porogen concentration results on lower gelation time as measured by photorheology. VP-POMaC with 40% porogen, but without TMPTA, resulted in a longer gelation time.

**Figure S6.**
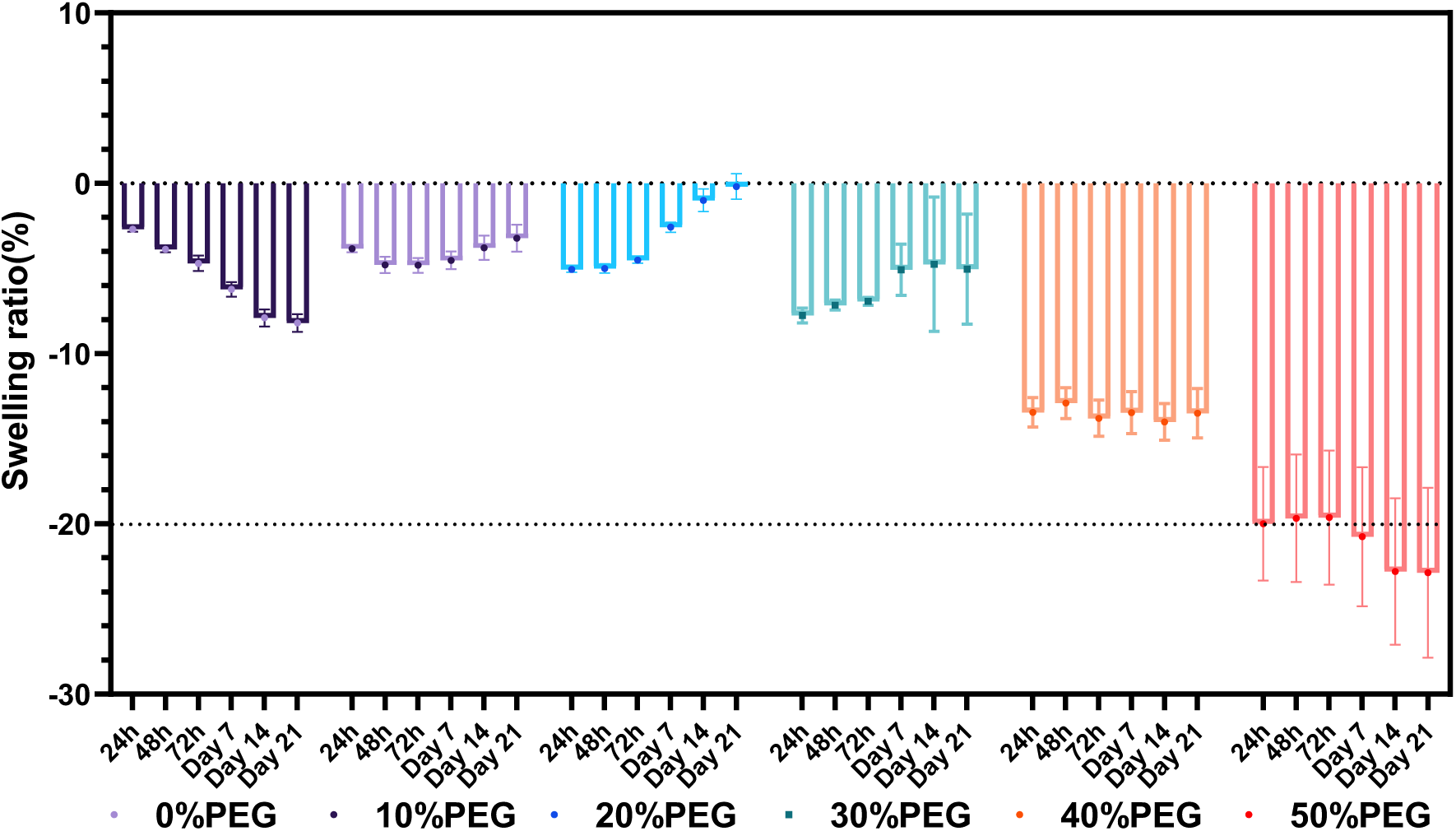
Swelling behaviour of VP-POMaC over 21 days in PBS for different porogen concentrations. Samples containing higher porogen concentrations exhibited greater shrinkage.

**Figure S7.**
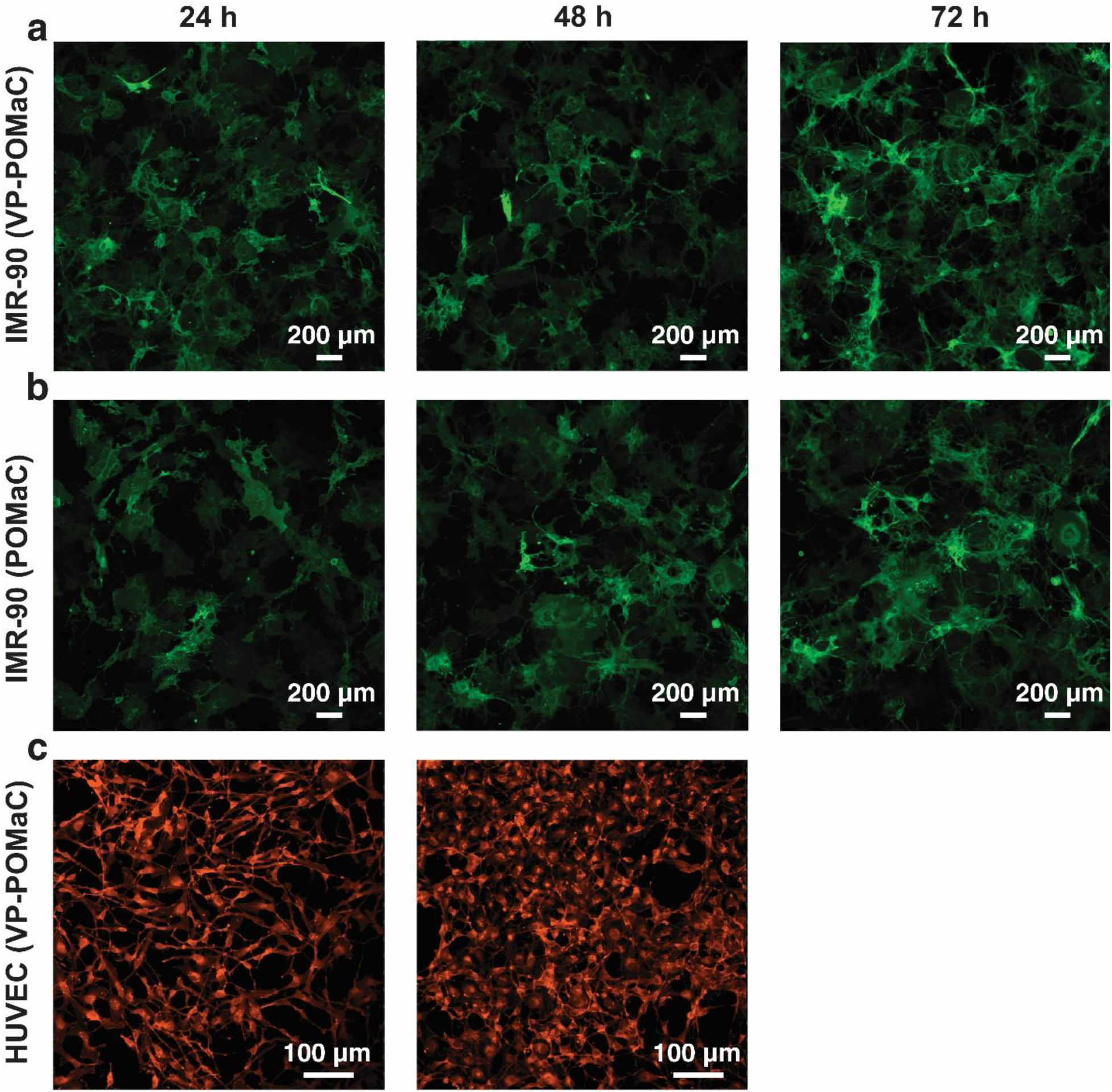
3D biocompatibility of VP-POMaC. **(a)** Fluorescence images of GFP labelled IMR- 90 human lung fibroblast cells following culture in a wellplate in presence of VP-POMaC at different time points. **(b)** Fluorescence images of GFP labelled IMR-90 human lung fibroblast cells following culture in a wellplate in presence of POMaC (previously published formulation) at different time points. We observed that cells co-cultured with VP-POMaC were morphologically indistinguishable compared to the benchmark POMaC formulation. **(c)** Fluorescence image of mCherry-labelled HUVEC cells following culture in a wellplate in the presence of VP-POMaC at different time points.

**Figure S8.**
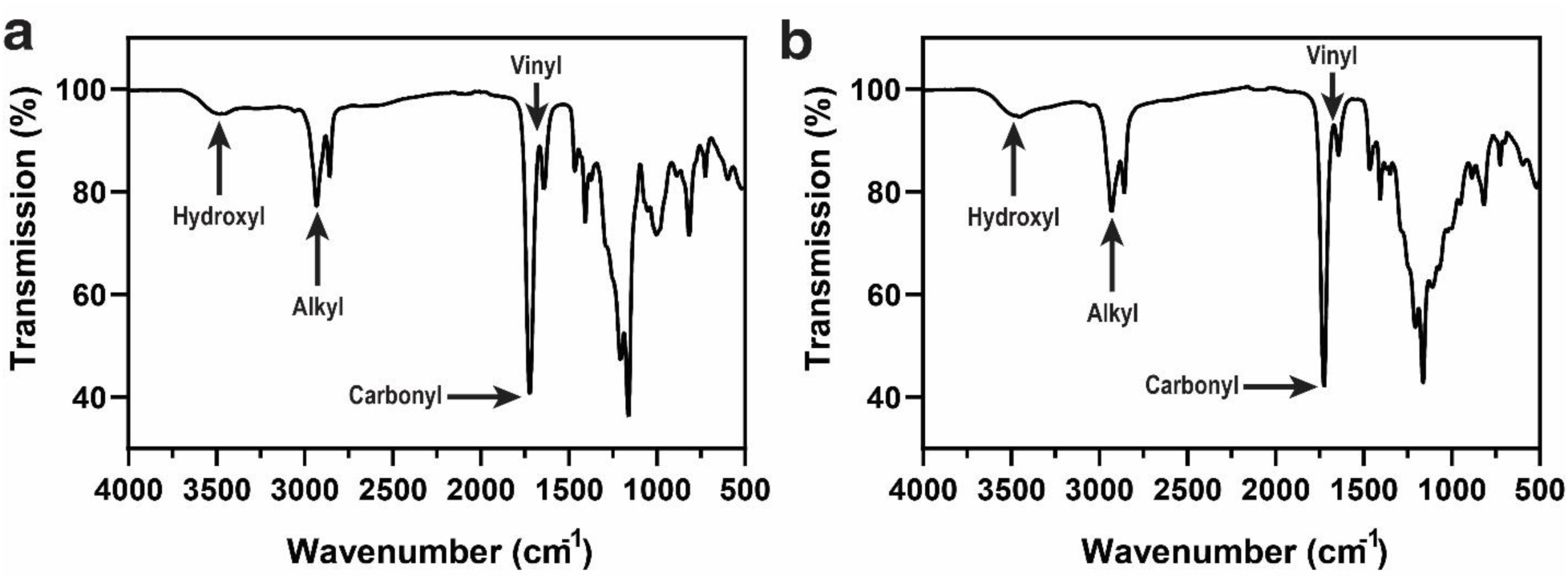
FT-IR characterization of POMaC. **(a)** FT-IR spectra of POMaC prepolymer**. (b)** FT-IR spectra of VP-POMaC film. FT-IR analysis verified the presence of various functional groups in the pre-polymer. FT-IR analysis of crosslinked VP-POMaC films reveals a decrease in the peak positioned at 1647 cm^−1^, attributed to the vinyl group from maleic anhydride.

**Table 1.** 3D printing parameters of VP-POMaC

| **Layer thickness (mm)** | **Base layer exposure time (s)** | **layer exposure time (s)** | **Rest time after retract (s)** | **Lifting distance (mm)** | **Retract speed (mm/min)** |
| --- | --- | --- | --- | --- | --- |
| 0.020 | 120 | 30-50 | 2 | 5 | 80 |

## References

1. Zhang, B., Korolj, A., Lai, B. F. L. & Radisic, M. Advances in organ-on-a-chip engineering. Nat. Rev. Mater. 1 (2018).

2. Ravanbakhsh, H. et al. Emerging Technologies in Multi-Material Bioprinting. Adv. Mater. 33, 2104730 (2021).

3. Ma, C., Peng, Y., Li, H. & Chen, W. Organ-on-a-chip: a new paradigm for drug development. Trends Pharmacol. Sci. 42, 119–133 (2021).

4. Nahak, B. K., Mishra, A., Preetam, S. & Tiwari, A. Advances in Organ-on-a-Chip Materials and Devices. ACS Appl. bio Mater. 5, 3576–3607 (2022).

5. Ghosh, U., Ning, S., Wang, Y. & Kong, Y. L. Addressing unmet clinical needs with 3D printing technologies. Adv. Healthc. Mater. 7, 1800417 (2018).

6. Li, T., Chang, J., Zhu, Y. & Wu, C. 3D printing of bioinspired biomaterials for tissue regeneration. Adv. Healthc. Mater. 9, 2000208 (2020).

7. Puza, F. & Lienkamp, K. 3D printing of polymer hydrogels—from basic techniques to programmable actuation. Adv. Funct. Mater. 32, 2205345 (2022).

8. Fuchs, S., Shariati, K. & Ma, M. Specialty tough hydrogels and their biomedical applications. Adv. Healthc. Mater. 9, 1901396 (2020).

9. Zhang, F. & King, M. W. Biodegradable polymers as the pivotal player in the design of tissue engineering scaffolds. Adv. Healthc. Mater. 9, 1901358 (2020).

10. Kirillova, A. et al. Fabrication of biomedical scaffolds using biodegradable polymers. Chem. Rev. 121, 11238–11304 (2021).

11. Zhang, B. et al. Biodegradable scaffold with built-in vasculature for organ-on-a-chip engineering and direct surgical anastomosis. Nat. Mater. 15, 669–678 (2016).

12. Tan, X. et al. Development of Biodegradable Osteopromotive Citrate-Based Bone Putty. Small 18, 2203003 (2022).

13. Davenport Huyer, L., et al. Tunable Bioelastomers: One-Pot Synthesis of Unsaturated Polyester Bioelastomer with Controllable Material Curing for Microscale Designs (Adv. Healthcare Mater. 16/2019). Adv. Healthc. Mater. 8, 1970064 (2019).

14. Wang, M., Xu, P. & Lei, B. Engineering multifunctional bioactive citrate-based biomaterials for tissue engineering. Bioact. Mater. 19, 511–537 (2023).

15. Tran, R. T. et al. Synthesis and characterization of a biodegradable elastomer featuring a dual crosslinking mechanism. Soft Matter 6, 2449–2461 (2010).

16. Zhang, B. et al. Microfabrication of AngioChip, a biodegradable polymer scaffold with microfluidic vasculature Microengineered biomimetic systems for organ-on-a. Nat. Protoc. 13, (2018).

17. Paunović, N. et al. Digital light 3D printing of customized bioresorbable airway stents with elastomeric properties. Sci. Adv. 7, eabe9499 (2021).

18. Liu, C. et al. High Throughput Omnidirectional Printing of Tubular Microstructures from Elastomeric Polymers. Adv. Healthc. Mater. 2201346 (2022).

19. Wales, D. J., Keshavarz, M., Howe, C. & Yeatman, E. 3D Printability Assessment of Poly (octamethylene maleate (anhydride) citrate) and Poly (ethylene glycol) Diacrylate Copolymers for Biomedical Applications. ACS Appl. Polym. Mater. 4, 5457–5470 (2022).

20. Lo Giudice, A., et al. Evaluation of the accuracy of orthodontic models prototyped with entry-level LCD-based 3D printers: A study using surface-based superimposition and deviation analysis. Clin. Oral Investig. 26, 303–312 (2022).

21. Katsuyama, Y. et al. A 3D-Printed, Freestanding Carbon Lattice for Sodium Ion Batteries. Small 18, 2202277 (2022).

22. Gallastegui, A. et al. Fast Visible-Light Photopolymerization in the Presence of Multiwalled Carbon Nanotubes: Toward 3D Printing Conducting Nanocomposites. ACS Macro Lett. 11, 303–309 (2022).

23. Kim, Y. T., Ahmadianyazdi, A. & Folch, A. A ‘print–pause–print’protocol for 3D printing microfluidics using multimaterial stereolithography. Nat. Protoc. 1–17 (2023).

24. Schüller-Ravoo, S., Feijen, J. & Grijpma, D. W. Preparation of flexible and elastic poly (trimethylene carbonate) structures by stereolithography. Macromol. Biosci. 11, 1662– 1671 (2011).

25. Kim, G.-T. et al. Cytotoxicity, Colour Stability and Dimensional Accuracy of 3D Printing Resin with Three Different Photoinitiators. Polymers (Basel). 14, 979 (2022).

26. Zeng, B. et al. Cytotoxic and cytocompatible comparison among seven photoinitiators- triggered polymers in different tissue cells. Toxicol. Vitr. 72, 105103 (2021).

27. Schneider, L. F. J., Cavalcante, L. M., Prahl, S. A., Pfeifer, C. S. & Ferracane, J. L. Curing efficiency of dental resin composites formulated with camphorquinone or trimethylbenzoyl-diphenyl-phosphine oxide. Dent. Mater. 28, 392–397 (2012).

28. Bhattacharjee, N., Parra-Cabrera, C., Kim, Y. T., Kuo, A. P. & Folch, A. Desktop- Stereolithography 3D-Printing of a Poly(dimethylsiloxane)-Based Material with Sylgard- 184 Properties. Adv. Mater. 30, 1–7 (2018).

29. Yan, Z. et al. Strengthening waterborne acrylic resin modified with trimethylolpropane triacrylate and compositing with carbon nanotubes for enhanced anticorrosion. Adv. Compos. Hybrid Mater. 5, 2116–2130 (2022).

30. Patel, D. K. et al. Highly stretchable and UV curable elastomers for digital light processing based 3D printing. Adv. Mater. 29, 1606000 (2017).

31. Harri, K. et al. Novel photo-curable polyurethane resin for stereolithography. RSC Adv. 6, 50706–50709 (2016).

32. Peng, S. et al. 3D printing mechanically robust and transparent polyurethane elastomers for stretchable electronic sensors. ACS Appl. Mater. Interfaces 12, 6479–6488 (2020).

33. Ji, Z. et al. 3D printing of photocuring elastomers with excellent mechanical strength and resilience. Macromol. Rapid Commun. 40, 1800873 (2019).

34. Zhao, Y. et al. A Platform for Generation of Chamber-Specific Cardiac Tissues and Disease Modeling. Cell 176, 913–927.e18 (2019).

35. Beauchamp, M. J., Nordin, G. P. & Woolley, A. T. Moving from millifluidic to truly microfluidic sub-100-μm cross-section 3D printed devices. Anal. Bioanal. Chem. 409, 4311–4319 (2017).

36. Parra-Cabrera, C., Achille, C., Kuhn, S. & Ameloot, R. 3D printing in chemical engineering and catalytic technology: structured catalysts, mixers and reactors. Chem. Soc. Rev. 47, 209–230 (2018).

37. Davenport Huyer, L., et al. Highly elastic and moldable polyester biomaterial for cardiac tissue engineering applications. ACS Biomater. Sci. Eng. 2, 780–788 (2016).

38. Tan, K., Cheng, S., Jugé, L. & Bilston, L. E. Characterising skeletal muscle under large strain using eccentric and Fourier Transform-rheology. J. Biomech. 48, 3788–3795 (2015).

39. Ramadan, S., Paul, N. & Naguib, H. E. Standardized static and dynamic evaluation of myocardial tissue properties. Biomed. Mater. 12, 25013 (2017).

40. Jacot, J. G., McCulloch, A. D. & Omens, J. H. Substrate stiffness affects the functional maturation of neonatal rat ventricular myocytes. Biophys. J. 95, 3479–3487 (2008).

41. Hrapko, M., Van Dommelen, J. A. W., Peters, G. W. M. & Wismans, J. The mechanical behaviour of brain tissue: large strain response and constitutive modelling. Biorheology 43, 623–636 (2006).

42. Ayyildiz, M., Cinoglu, S. & Basdogan, C. Effect of normal compression on the shear modulus of soft tissue in rheological measurements. J. Mech. Behav. Biomed. Mater. 49, 235–243 (2015).

43. Zhan, Y., Fu, W., Xing, Y., Ma, X. & Chen, C. Advances in versatile anti-swelling polymer hydrogels. Mater. Sci. Eng. C 127, 112208 (2021).

44. Boutry, C. M., Kaizawa, Y., Schroeder, B. C., Chortos, A. & Legrand, A. A stretchable and biodegradable strain and pressure sensor for orthopaedic application. (2018).

45. Montgomery, M., et al. Method for the Fabrication of Elastomeric Polyester Scaffolds for Tissue Engineering and Minimally Invasive Delivery. ACS Biomater. Sci. Eng. 4, 3691–3703 (2018).

46. Zhang, B. et al. Microfabrication of AngioChip, a biodegradable polymer scaffold with microfluidic vasculature. Nat. Protoc. 13, 1793 (2018).

47. Hayer, A. et al. Engulfed cadherin fingers are polarized junctional structures between collectively migrating endothelial cells. Nat. Cell Biol. 18, 1311–1323 (2016).

